# Optogenetic Control of cAMP Levels and HCN Channels: Implications in Cardiac Physiology and Parkinson’s Disease

**DOI:** 10.64898/2026.08.13.744738

**Authors:** Run-Zhou Yang, Dian-Dian Wang, Dan-Hua Liu, Pei-Pei Liu, Shu-Ang Li, Jian-Sheng Kang

## Abstract

Cyclic adenosine monophosphate (cAMP) is a second messenger that regulates various cellular processes, including the activity of hyperpolarization-activated channels (HCN), which are implicated in cardiac physiology and neurodegenerative diseases such as Parkinson’s disease (PD). In this study, we used a photoactivated adenylyl cyclase (PAC) S27A mutant to optogenetically control intracellular cAMP levels. We demonstrated that light-induced elevation of cAMP activated HCN4 channels, leading to increased beating rates in cardiomyocytes. Unilateral expression of PAC(S27A) in the *substantia nigra pars compacta* of mice induced rotation behavior upon light stimulation, which could be attenuated by HCN inhibitors. Furthermore, PAC(S27A) activation partially recovered motor deficits in a 1-methyl-4-phenyl-1,2,3,6-tetrahydropyridine (MPTP)-induced PD mouse model, accompanied by increased HCN2 channel expression in ipsilateral basal ganglia. Our findings highlight the potential of using optogenetics to modulate cAMP and HCN channel activity for the treatment of cardiac and neurological disorders.

## Introduction

Cyclic adenosine monophosphate (cAMP) is a second messenger that is widely distributed from prokaryotes to humans and plays an important role in regulating cellular transcription, translation, metabolism, and protein transport(*1*). In different cells, cAMP affects physiological activities by acting on different target proteins, including PKA, EPAC, CNGC, and HCN(*2*, *3*). cAMP can affect the opening of hyperpolarization-activated channels (HCN) channels by binding to the cAMP-binding domain on the HCN channel(*4*). For example, in the treatment of cardiac arrest, adrenaline injection activates adenylyl cyclase, increasing the concentration of cAMP in the myocardium to bind to HCN4 channels, increasing their permeability and triggering faster heartbeats(*5*). cAMP has a short half-life in cells and is generally hydrolyzed by intracellular phosphodiesterases (PDEs) to produce AMP after exerting its effects to prevent sustained cell activation(*6*). In the nervous system, cAMP can affect axonal and dendritic growth, dendritic spine formation, and synaptic plasticity(*7*, *8*).

Hyperpolarization-activated channels (HCN) are involved in various pathological processes such as Alzheimer’s disease and Parkinson’s disease(*9*, *10*). In Alzheimer’s disease monkeys, decreased expression of HCN1 in the temporal lobe is associated with increased Aβ production in the olfactory cortex (*10*). In Parkinson’s disease, HCN channels are related to freezing behavior and synchronous oscillations and high-frequency bursts in the *substantia nigra pars compacta* (SNC) (*10*). In PD mice, the Ih current of dopaminergic neurons in the SNC is decreased before the onset of PD symptoms (*11*). HCN channels play a role in sleep, where they are related to the production of low-frequency high-amplitude EEG delta waves in non-rapid eye movement sleep(*12*). They also involve in anesthesia and anesthetics could reduce the activities of HCN channels(*13*, *14*).

The activity of HCN channels depends on cAMP levels(*15*). Regulating cAMP levels can affect HCN channel activity and influence cell electrophysiology. There are transmembrane ACs (AC1-AC9) activated by G proteins, and soluble ACs (sACs) activated by bicarbonate(*16*). sACs also exist in nuclei and mitochondria. Mitochondrial sAC can act as a metabolic sensor(*17*). Optogenetics are playing emerging roles in controlling cAMP signaling. There are two ways to use light to increase intracellular cAMP: 1) Import an exogenous light-sensitive G protein-coupled receptor (GPCR) to activate endogenous ACs; 2) Import an exogenous light-sensitive AC to directly produce cAMP from ATP. OptoXR is a class of optogenetic tools developed in 2009 to regulate intracellular signaling using light. It includes opto-α1AR and opto-β2AR(*18*). Opto-β2AR combines the light sensitivity of rhodopsin and the G protein-activating ability of the β2-adrenergic receptor. Under light, it activates G proteins and increases cAMP. Photoactivated adenylyl cyclase (PAC) is a light-sensitive AC discovered in *Euglena gracilis*(*19*). It has been used to treat infertility in mice(*20*). Recently, PAC is also used to activate thermogenesis in brown adipocytes which provides potential therapies that target obesity-related disorders(*21*). Wild-type PAC has some activity in the dark. Mutagenesis and screening identified mutants (S27A, H226W, F198Y, K197A) with lower dark activity and higher light-activated AC activity. Among them, S27A and H226W showed the biggest increases(*22*). Compared to wild-type PAC, the mutants enable more precise regulation of cAMP using light due to their lower background activity in the dark.

In this study, PAC S27A mutant was used to control the intracellular cAMP level. Elevated cAMP by light could activate HCN4 channel activities and cardiomyocytes’ beating. Furthermore, unilateral PAC S27A expression in *substantia nigra pars compact* induced mice rotation behavior upon light stimulation, which can be attenuated by HCN inhibitor. PAC S27A activation partial recovered the mice motor deficit induced by MPTP, accompanied by an increased expression of HCN2 channel in ipsilateral basal ganglia.

## Results

Overexpression of PAC(S27A) in 293t cells leads to an increase of intracellular cAMP levels under light exposure as indicated by cAMP indicators H74 (Figure 1A-B). Co-expression of HCN4 and PAC(S27A) in 293t cells under light exposure results in an inward current measured by patch clamping (Figure 1C-D), suggesting that light activates PAC(S27A) to open HCN4 channels. Illumination with different wavelengths indicates that blue light (488 nm) produces a larger photocurrent than green light (540 nm) and violet light (405 nm) (Figure 1D).

**Figure 1.**
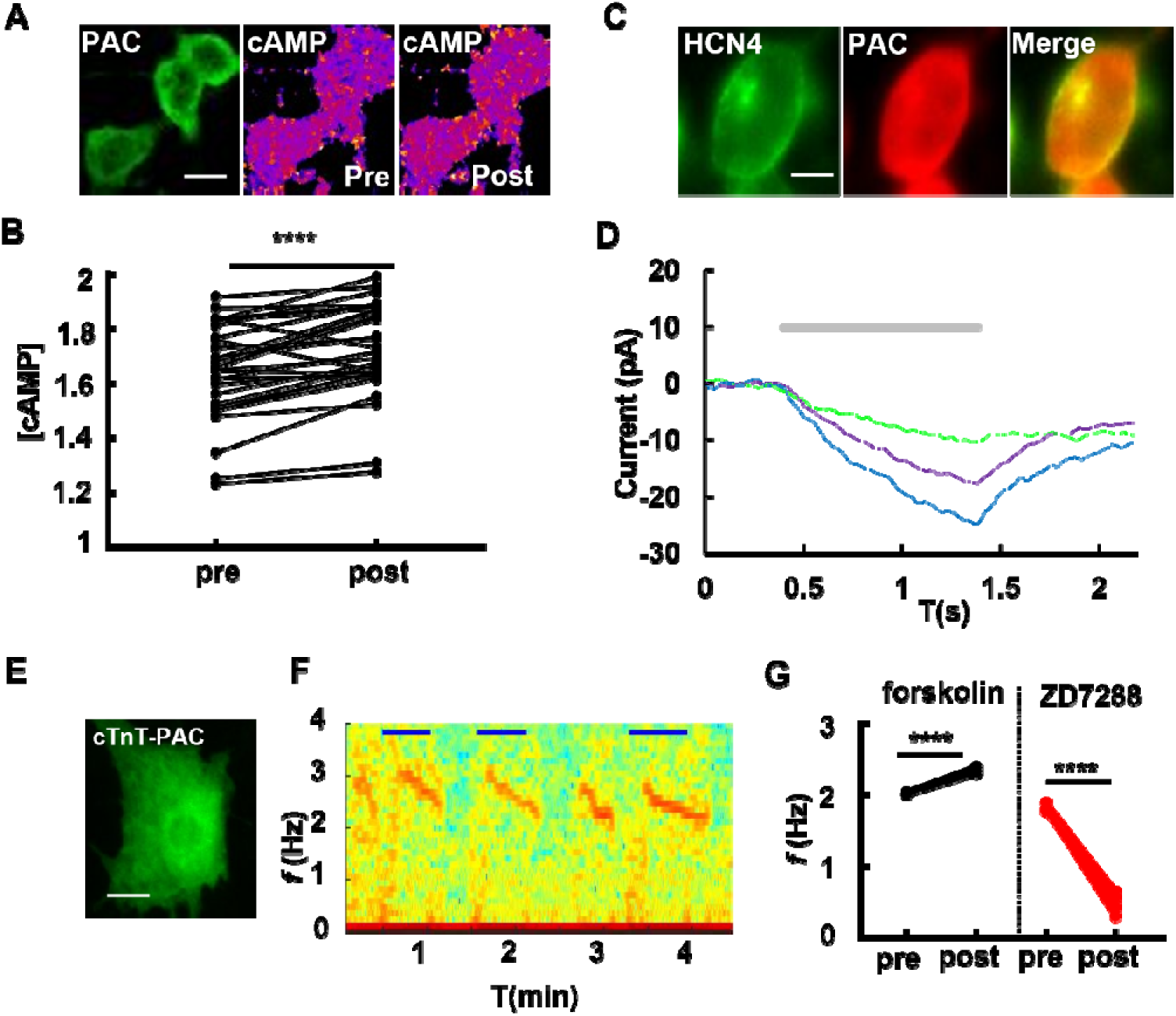
Characteristics of photoactivated adenylyl cyclase (PAC). **(A)** Expression of PAC(S32A) and cAMP indicator H74 in HEK293 cells, showing changes in fluorescent ratio before (left) and after light stimulation (right). Scale bar, 5 μm. **(B)** Changes in cAMP levels in PAC-expressing cells before and after light stimulation measured with H74 (paired *t-test*, *p* < 0.0001). **(C)** Co-expression of cAMP-sensitive HCN4 channels with PAC in HEK293t cells. Scale bar, 5 μm. **(D)** Photocurrents of HEK293t cells expressing HCN4 channels and PAC under different wavelengths of light (405 nm, purple; 488 nm, blue; 532 nm, green) measured by patch clamp. **(E)** Expression of PAC in cardiomyocyte. Scale bar, 5 μm. **(F)** Frequency map of beating frequency of PAC-expressing cardiomyocytes during light stimulation measured with video analysis. Blue bar represents 488 nm light. **(G)** Changes in intracellular Ca^2+^ frequency in cardiac myocytes after treatment with adenylyl cyclase activator forskolin and HCN channel inhibitor ZD7288 (paired *t-test*, *p* < 0.0001).

To investigate the effect of activation on cardiomyocytes, PAC(S27A) was expressed under the control of the cardiac-specific promoter, cTnT (Figure 1E). Upon light stimulation, an increment was observed in the beating rate of the cardiomyocytes (Figure 1F). Intriguingly, we observed a delay in beating upon illumination. The beating rate started to increase approximately 15 seconds after the light onset and began to decrease almost 10 seconds after light offset (Figure 1F). We hypothesize that cAMP plays a crucial role in controlling cardiomyocyte beating by affecting HCN channel activity, as PAC(S27A) produces more cAMP under light. To assess the roles of cAMP and HCN in cardiomyocytes, an AC agonist, forskolin, and an HCN inhibitor, ZD7288, were added to cultured cardiomyocytes. As expected, forskolin increased the beating rate while ZD7288 decreased it (Figure 1G).

To further explore the function of PAC(S27A) in neuronal systems, we expressed it unilaterally in the *substantia nigra pars compacta* (SNC) via AAV delivery (Figure 2A). Four weeks after virus injection, mice were placed in an open-field box for light stimulation and behavior recording (Figure 2B). An ipsilateral rotation behavior was observed upon exposure to 488 nm light (Figure 2E-F), which disappeared when the light stopped, indicating rapid hydrolysis of cAMP produced by PAC *in vivo*. To confirm that the light-induced rotation behavior was not due to the effect of light on animal behavior, we changed the wavelength of the light. As the wavelength increased, the effect of light-induced rotation behavior decreased (Figure 3), consistent with the PAC light sensitivity observed by patch clamping (Figure 1D). Injection of HCN channel inhibitors ZD7288 or zatebradine into the SNC before PAC activation did not result in significant light-induced rotational behavior (Figure 1G-I), indicating that the cAMP produced by PAC affects the excitability of substantia nigra neurons through HCN channel activation. Decreased calcium levels were observed in the SNC brain region after the addition of HCN channel inhibitor zatebradine (Figure 2J), indicating that the SNC brain region relies on HCN channels to maintain excitability in its physiological state.

**Figure 2.**
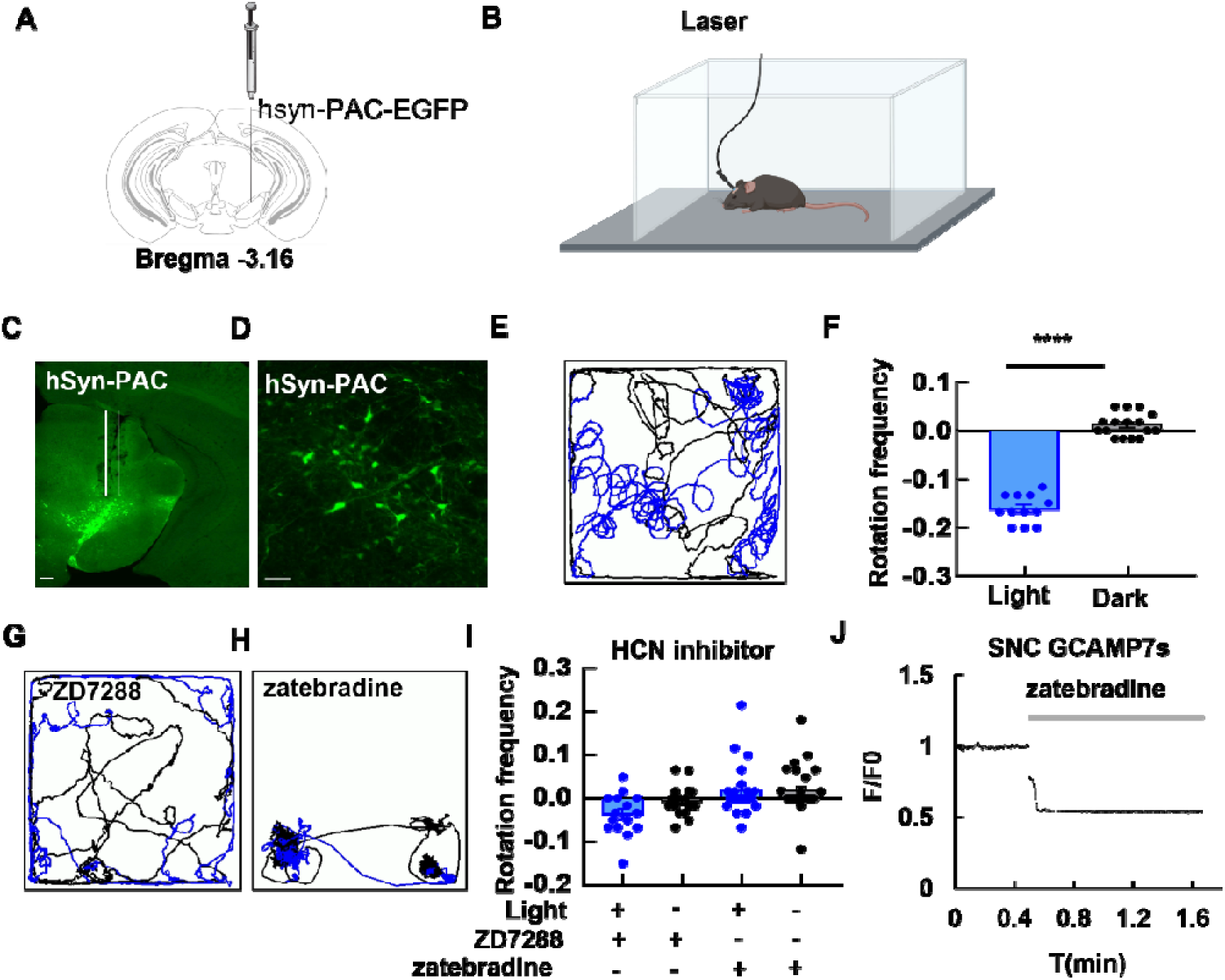
Light-induced rotation behavior in mice with PAC (S27A) injected into the substantia nigra. **(A)** A diagram showing injection of virus into one side of the substantia nigra pars compacta (SNC, ML -1.13, AP -3.16, DV -4.4) of the brain. **(B)** A diagram showing behavior testing upon light stimulation in an open field box. **(C)** Brain slice of a mouse injected with hSyn-PAC-EGFP. **(D)** A zoomed-in view of brain slice in (C). **(E)** Movement trajectory of a mouse injected with hSyn-PAC-EGFP during light stimulation (473 nm, 10 mW). **(F)** Rotation frequency of mice injected with hSyn-PAC-EGFP before and after light stimulation, with ipsilateral rotation as negative values and contralateral rotation as positive values. Significant ipsilateral rotation was observed after light stimulation (*t-test*, *p* < 0.001). **(G)** Movement trajectory after injection of HCN channel inhibitor ZD7288 into the substantia nigra of mice expressing hSyn-PAC-EGFP. The time period stimulated with light was indicated as blue. **(H)** Movement trajectory after injection of HCN channel inhibitor zatebradine into the substantia nigra of mice expressing hSyn-PAC-EGFP. The time period stimulated with light was indicated as blue. **(I)** Rotation frequency of mice injected with hSyn-PAC-EGFP before and after HCN channel inhibitor ZD7288 or zatebradine injection into the substantia nigra and light stimulation. **(J)** Changes in Ca^2+^ levels in the substantia nigra region after injection of HCN channel inhibitor zatebradine into the substantia nigra.

**Figure 3.**
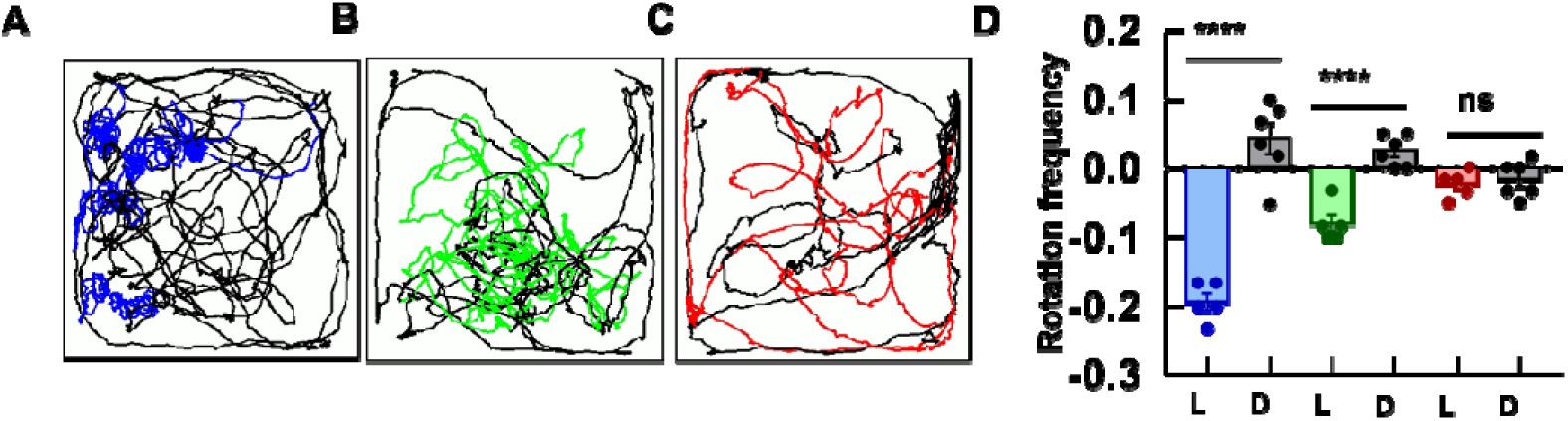
Activation of PAC by light of different wavelengths induce different extent of rotation behavior. **(A)** Movement trajectory of a mouse injected with hSyn-PAC-EGFP in the substantia nigra with blue light stimulation (473nm, 10mW). **(B)** Movement trajectory of a mouse injected with hSyn-PAC-EGFP in the substantia nigra during green light stimulation (532nm, 10mW). **(C)** Movement trajectory of a mouse injected with hSyn-PAC-EGFP in the substantia nigra during red light stimulation (633nm, 10mW). **(D)** Rotation frequency of mice injected with hSyn-PAC-EGFP under different wavelength light stimulation, with ipsilateral rotation as negative values and contralateral rotation as positive values (blue bar: blue light, green bar: green light, red bar: red light, gray bar: dark). Significant differences were observed under blue light (*t-test*, p < 0.001), while there was no significant difference under red light (*t-test*, p > 0.05).

To further investigate the effects of PAC(S27A) in relieving PD-like symptoms, a MPTP induced PD mouse model was constructed. By Intraperitoneal injection with MPTP, mouse EEG was monitored (Figure 4A). A theta wave was observed after MPTP injection, accompanied by the decreasing of calcium level and frequency (Figure 4B-C). After MPTP injection, light was delivered to the side of SNC with PAC(S27A) injected. Light stimulation could still induce rotation behavior after MPTP treatment (Figure 4D-E), although the frequency is largely decreased compared with that under normal conditions. The cAMP level in SNC was dramatically decreased upon MPTP treatment, as measured by flamindo2(*23*) (Figure 4G), while the ATP level indicated by iATP in SNC was relatively constant (Figure 4F). To the identify the mechanism of the protective effect of PAC(S27A) on PD, we detected the expression of various HCN channels in the brain using immunofluorescence and found that the expression of HCN2 protein in the striatum on the activated side of light exposure was significantly increased (Figure 5), while the expression of other HCN channels remained unchanged.

**Figure 4.**
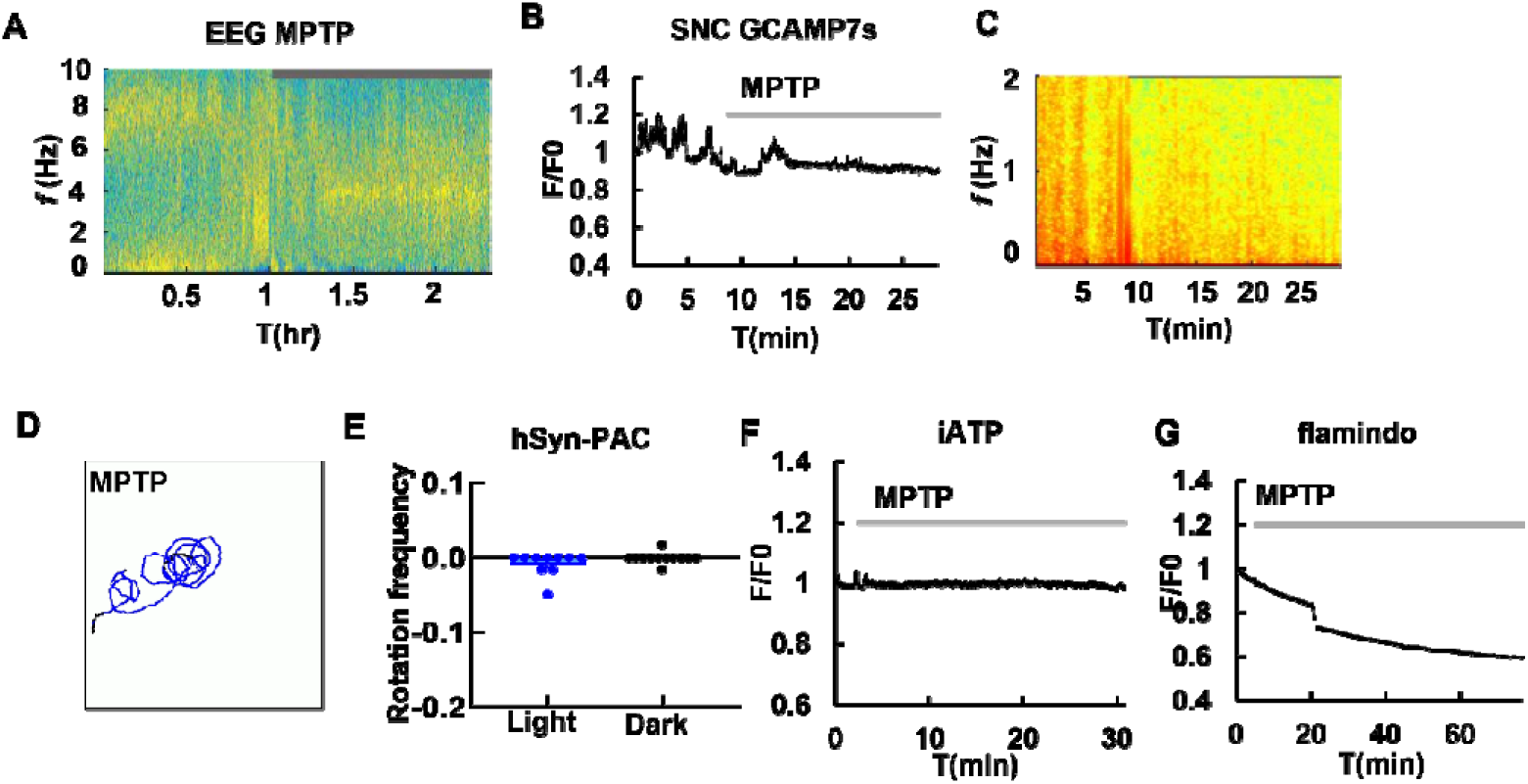
Effects of MPTP treatment and activation of PAC in mice. **(A)** Electroencephalogram (EEG) of mice before and after intraperitoneal injection of MPTP. **(B)** Ca^2+^ levels in substantia nigra compacta neurons before and after intraperitoneal injection of MPTP. **(C)** Frequency of Ca^2+^ oscillation in substantia nigra compacta neurons before and after intraperitoneal injection of MPTP. **(D)** Movement trajectory of a mouse expressing hSyn-PAC injected into the substantia nigra during blue light stimulation (473 nm, 10 mW) after intraperitoneal injection of MPTP. **(E)** Rotation frequency of mice injected with hSyn-PAC-EGFP under light stimulation after MPTP treatment, with ipsilateral rotation as negative values and contralateral rotation as positive values. **(F)** Intracellular ATP levels in substantia nigra compacta before and after MPTP injection. **(G)** Intracellular cAMP levels in substantia nigra compacta before and after MPTP injection.

**Figure 5.**
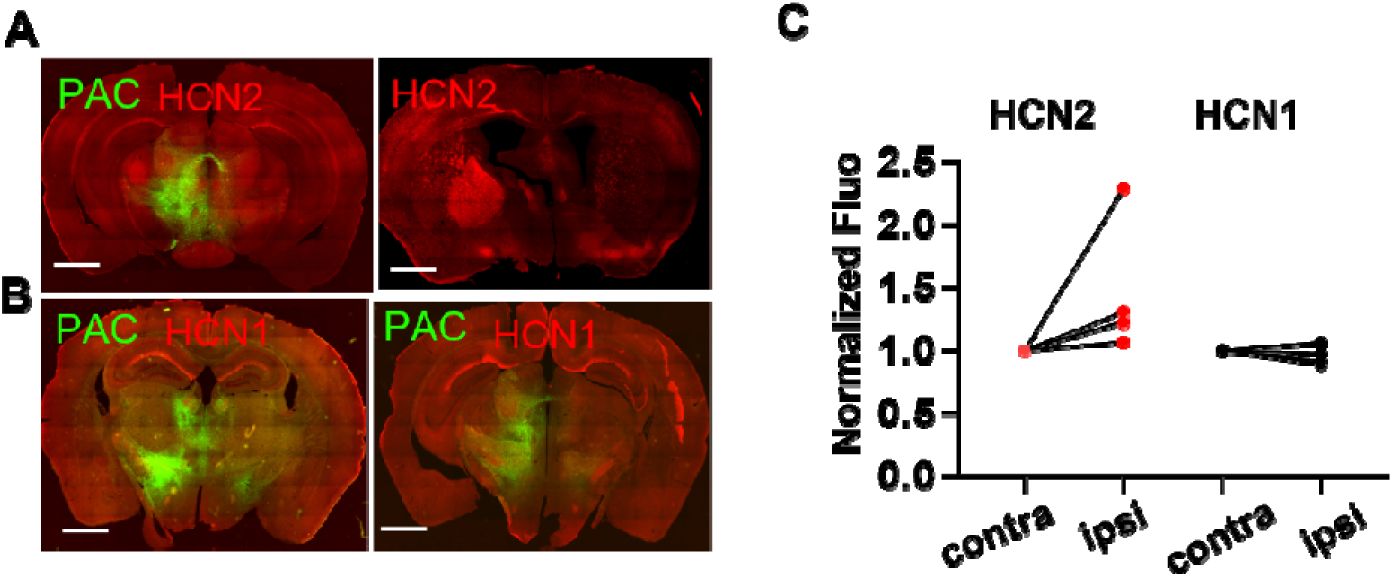
Activation of PAC leads to increased expression of HCN2 on the same side. **(A)** HCN2 staining in substantia nigra of mice expressing hSyn-PAC after light activation. **(B)** HCN1 staining in substantia nigra of mice expressing hSyn-PAC after light activation. **(C)** Comparison of HCN2 and HCN1 staining in mice expressing hSyn-PAC after light activation, comparing fluorescence on the contralateral and ipsilateral sides.

## Discussion

cAMP is playing emerging roles in cardiovascular diseases(*24*) and neuronal disorders (*25*). Traditional optogenetic tools, such as hChR2 and NpHR, are based on ion flux that influence the action potential. Modulating cAMP levels by optogenetics provides a new avenue to regulate cell metabolism. Previous studies demonstrates that photoactivation of PAC during tetanic stimulation enabled synaptic potentiation(*26*). In our results, we demonstrate that PAC(S27A) was able to induce HCN4 channels opening and accelerating cardiomyocytes’ beating rate. The HCN4 current induced by PAC activation exhibited a time-dependent manner, which might be contributed to the accumulating cAMP produced by PAC. In addition, the cardiomyocytes began to beat faster after illumination for a few seconds, rather than beat immediately upon light stimulation. This indicates that there exists a threshold level of cAMP before eliciting physiological reactions. Previous studies showed that PKA is much less sensitive to cAMP in cells than in vitro with an activation threshold 20 fold higher in vivo, which is contributed to the compartmentation of cAMP (*27*). It’s possible that the as cAMP produced from PAC(S27A) is generally distributed in the cytosol, the cAMP should accumulate to a level that could activate downstream effectors. Compartment of PAC may improve the sensitivity, as the previous reported a plasma membrane-anchored PAC (PACmn) with improved light sensitivity and decreased dark activity(*22*). Downstream effectors of cAMP include PKA and HCN. PKA regulates calcium cycling and myofibril contractility by phosphorylating different proteins including L-type calcium channel, RyR, phospholamban, Troponin I, and MyBP-C(*8*). HCN4 and HCN2 are modulated by cAMP, while HCN1 and HCN3 are less sensitive (*10*).

Rotation behavior has been observed in unilateral injection of GABA agonists, glutamate antagonists into substantia nigra(*28–32*). PAC(S27A) unilateral injection resulted in an ipsilateral rotation, which is in line with previous reports conducted with drugs. Pretreatment with HCN channel inhibitors ZD7288 weaken the effects on rotation. In mouse treated with zatebradine, a contralateral turning was observed despite of light stimulation, accompanied by a decrease of calcium level. These results reveal an important role of HCN channels in substantia nigra in controlling locomotion. Previous studies showed that spontaneous firing activity of dopaminergic neurons can be blocked by ZD7288(*33*). Injecting ZD7288 unilaterally into the SNC of adult rats significantly elevated the number of rotations induced by apomorphine, while decreasing the immunofluorescence intensity of tyrosine hydroxylase(*34*).

MPTP application resulted in an increase of theta (2–7 Hz)-frequency bands in our results. The occurrence of theta activity has been associated with a paroxysmal escalation of freezing behavior in patients with Parkinson’s disease(*35*, *36*). HCN channels are reported to be involved in generating theta oscillation(*37*). Exposure to MPP^+^ leads to the occurrence of damage to the SNc within 0.5 - 2 hours (*38*). Although the Ih current was reduced by MPP^+^, MPP^+^ inhibits pacemaker activity of SNC neurons by activation of ATP-sensitive potassium (KATP) rather than directly block Ih(*38*). PAC(S27A) activation results in a rotation behavior after MPTP treatment, indicating that the by increasing cAMP level could partially improve the motor deficits induced by MPTP. A significant decrease of cAMP upon MPTP was observed, while intracellular ATP remains constant, which indicated that cAMP decrease is an early event during MPTP treatment, while intracellular ATP decreased at a later stage when energy was depleted.

The decrease in HCN channel expression in dopaminergic neurons of the SNc is one of the initial physiological alterations linked to Parkinson’s disease in MitoPark mice(*11*). Within the basal ganglia, HCN2 and HCN4 are relatively more highly expressed than HCN3 in the SNc(*10*). Meanwhile, HCN2 serves as the primary isoform in the GP, whereas both HCN2 and HCN3 are more prominently expressed in the STN(*10*). In rats treated with 6-hydroxydopamine (6-OHDA), the expression of HCN2 in the external globus pallidus (GPe) decreases(*39*), while the HCN3 expression was selectively upregulated in the GPi(*40*). After PAC(S27A) activation, we observed an increased expression of HCN2 channels in the globus pallidus of ipsilateral side, indicating that cAMP elevation induced by PAC(S27A) could also regulate HCN channels at transcriptional levels.

Overall, this study highlights the potential of using PAC(S27A) to control the intracellular cAMP level and regulate HCN channel activity, which could have implications for the development of new therapies for various diseases, such as cardiovascular disease and neurological diseases.

## Material and Methods

### Plasmid construct

PAC (NCBI Accession ID: ADC33126) was synthesized according to human codon usage. The H74 cAMP sensor was gifted from Kees Jalink, while the sequences of iATP and flamido2 were obtained from Addgene (Addgene ID: 102550, 73938). The mouse HCN4 channel was cloned from mouse heart mRNA (NCBI Accession ID: NM_001081192.3) and fused with EGFP in the pEGFPN1 vector. To enable expression in cardiomyocytes, the cTnT promoter (-1 to -589) was amplified from mouse genome DNA. For neuronal expression, the hSyn promoter (-1 to -448) was amplified from plasmid. For AAV virus packaging, genes with promoters were cloned into pAAV-MCS vectors. All constructs were verified by sequencing (Genewiz).

### Cell culture and transfection

The HEK293t and Hela cell lines were cultured in DMEM (Gibco) supplemented with 10% fetal bovine serum. The cells were transfected using the calcium phosphate precipitation method and were incubated at 37°C in a humidified atmosphere containing 5% CO_2_ and 95% air.

### Living cell imaging and light stimulation

Hela cells transfected with H74 and PAC-mKate2 were imaged using a two-photon imaging system (Leica 980). H74 was excited with a two-photon laser set at a wavelength of 820 nm, and the emission was collected in the 450-490 nm range (CFP channel) and 520-550 nm range (YFP channel). The cAMP level was measured as the ratio of the two emission channels. For light stimulation, cells were exposed to a halogen lamp filtered by a FITC excitation filter (467-498 nm) for 5 minutes. Images were acquired both before and after light illumination.

### Electrophysiology

Whole-cell patch clamp recordings were conducted on HEK293t cells at room temperature using a customized opto-electro system. The setup included an Axopatch 700B amplifier manufactured by Molecular Devices, a digitizer (Digidata 1440A, Molecular Devices), a monochromator (Optoscan, Cairn Research Ltd., UK), and an imaging system made by Olympus. The devices were controlled by customized Micro-Manager software. A sampling rate of 10 kHz was utilized to acquire the data. Micropipettes were pulled from filamented glass capillaries (Sutter Instrument, BF150-86-10) using a micropipette puller (Sutter Instrument, P1000) to achieve a tip resistance of 4-6 MΩ. The micropipette was filled with intracellular buffer solution (composed of potassium gluconate 120 mM, KCl 3 mM, HEPES 10 mM, NaCl 8 mM, CaCl2 0.5 mM EGTA 5 mM, ATP-Mg 2 mM, GTP 0.3 mM, pH 7.2) and positioned using a micromanipulator (Sutter, MP285). To record light-induced photocurrents, HEK293t cells were placed in Tyrode’s buffer at room temperature and a customized protocol was used to generate light pulses with durations of 1 second and intervals of 1 second.

### Primary cardiomyocyte culture

Postnatal day 0 (P0) C57BL mice hearts were dissected and cut into small pieces, which were then washed with HBSS. To digest the tissues, trypsin (Sigma-Aldrich) was applied for 5 minutes, followed by collagenase II (Sigma-Aldrich) for an additional 30 minutes. The cells were further dissociated using fire-polished pipettes before being plated onto matrix gel-coated coverslips (Glasswarenfabrik Karl Hecht, Germany, 12 mm). Cardiomyocytes were cultured in plating medium and fed twice a week. Within 24 hours, the cardiomyocytes began to beat spontaneously. For AAV transduction, the virus was added to the medium at DIV2-3, and the cells were imaged at DIV7-8.

### Virus preparation and transduction

The animals were transfected with AAV DJ serotype virus. The AAV virus was generated by co-transfecting gene plasmid, capsid (pAAV-DJ), and helper plasmids (pHelper) into 293t cells using the calcium phosphate precipitation method. After 48 hours post-transfection, viruses were obtained from cell pellets through four cycles of freeze-thaw methods.

### Video-based cardiomyocytes beating analysis

According to the Nyquist sampling rule, the imaging sampling frequency should be more than twice the frequency of cardiomyocyte beating. In this study, bright field images were captured at a rate of 8Hz using an EMCCD camera (Photometric Evolve).

The acquired video was analyzed with ImageJ (NIH). Firstly, the video was converted to 8-bit and a line was drawn on it to track the movement of cells. Subsequently, a Kymograph was generated using the MultipleKymograph plugin. By drawing a straight line across the time-axis of the kymograph, a plot of grayscale value versus time was obtained using the “Plot Profile” command. The grayscale-value data was saved and used for spectrogram analysis. Finally, the MATLAB spectrogram function was employed to generate the spectrogram.

### Animals

Wild type mice (C57/BL 6J) were acquired from Charles River Laboratories. The mice were kept under a 12-hour light/dark cycle and had unlimited access to food and water, with the exception during behavioral experiments. All procedures were in accordance with protocols that were approved by the Animal Ethics Committee of Zhengzhou University.

### Mouse stereotactic injection

Adult mice weighing 24-26g were anesthetized with 3% pentobarbital sodium (intraperitoneal injection, 30 mg/kg) and fixed in a stereotaxic apparatus (RWD Instruments, Shenzhen, China). The skull was exposed and treated with hydrogen peroxide to visualize the bregma and lambda site. Glass pipettes (3.5# Drummond, USA) were pulled using a programmable micropipette puller (Sutter Instruments, USA). The target site was SNC (AP, -3.16 mm, ML ±-1.13 mm, DV, -4.4 mm). A drill was used to create a hole on the skull for virus injection. 500-1000 nL of virus was injected at a flow rate of 100 nl/min using an injection pump (KD Scientific, USA). Following injection, the pipette remained in place for about 5 minutes before being withdrawn. Optical fibers (diameter: 200 μm, numerical aperture: 0.37, length: 5 mm, RWD Instruments, Shenzhen, China) were implanted and secured with dental cement (Yamahachi, Japan) for light stimulation. For fiber photometry, optical fibers with a larger diameter (diameter: 400 μm, numerical aperture: 0.37, length: 5 mm, RWD Instruments, Shenzhen, China) were utilized. For simultaneous drug delivery and fiber imaging, a combined fiber with cannula (diameter: 200 μm, numerical aperture: 0.37, length: 5 mm, Kedoubc, China) was implanted. The injection and fiber implantation sites were confirmed in tissue sections.

### Light stimulation

Mice were placed in an open-field box for light stimulation and behavioral recording. The transplanted fiber of the mouse was connected to an optic cable through a rotary joint to the lasers (Dream Lasers, Shanghai). An Arduino-controlled customized device was used to generate laser pulses and record the video of the mouse’s behavior during light stimulation. For PAC stimulation, 473 nm, 532 nm, and 633 nm lasers were used to generate light pulses that lasted for 2 minutes each with 1-minute intervals. The laser intensity was measured to be approximately 10 mW using a laser power meter (Sanwa, Japan).

### Cannula drug delivery

The mouse’s cannula was linked to a PE tube and syringe controlled by a microinjection pump. A dose of 5 μl of the drug zatebradine (100 μM) was injected into the cannula at a rate of 2 μl/min.

### MPTP injection and EEG measurement

The mice were intraperitoneally injected with MPTP at a dosage of 20mg/kg. EEG measurements were taken using a bio-signal recording system (Medusa, HK, China). The mice were anesthetized with 4% chloral hydrate (intraperitoneal injection, 400 mg/kg) and immobilized in a stereotaxic apparatus (RWD Instruments, Shenzhen, China). The skull was exposed, and the EEG electrode was fixed to the mouse’s skull by attaching it to four cranial pins. Additionally, EMG wires were inserted into the neck muscles. For EEG recording, the EEG electrode was connected to the amplifier, which in turn was linked to the recording system and computer.

### Behavior analysis

The analysis of mouse behavior was performed using customized Python scripts based on the Deeplabcut toolbox(*41*). Labeled images were trained using a pretrained resnet50 network over 100,000 iterations. The models were evaluated and fine-tuned to achieve an accuracy of at least 95%. After training, the models were exported and used for further behavior analysis. To assess the rotation of the mouse, the angular acceleration of the angle between the vector formed by the head and tail and the horizontal vector was measured as the rotation frequency.

### Brain slices

The mouse was anesthetized with an intraperitoneal injection of 4% chloral hydrate (400 mg/kg) and subsequently perfused with normal saline and 4% PFA for tissue fixation. The brain was then carefully extracted from the skull and placed in a 40% sucrose solution for dehydration before being embedded in OCT compound (SAKURA Tissue-Tek, Japan) for frozen sectioning. Slices were cut to a thickness of 30 μm using a freezing microtome (Leica, CM 3050S, Germany) and visualized using a fluorescence microscope (Olympus IX83, Japan).

### Fiber photometry

The fiber transplanted into the mouse was connected to an optic cable that led to a fiber photometry detector (Thinker, China). To capture images of iATP or Flamingo2, the proteins were excited with 488 nm light, and their emissions were collected in the 490-540 nm range. Adjustments were made to the light intensity and fluorescence signal gains to prevent rapid bleaching of the fluorescence.

### Immunostaining

For immunostaining of brain slices, the slices were exposed to 0.1% Triton X-100 to increase permeability for the antibody. After blocking with 1% BSA, the slices were incubated with primary antibody (HCN2, 1:100, rabbit, Sigma-Aldrich) overnight.

The next day, the primary antibody was washed off, and the slices were incubated with the secondary antibody (Alexa Fluor 555, anti-rabbit, 1:1000, Thermo, USA). Finally, the slices were imaged using a fluorescence microscope (Olympus IX83, Japan).

## Author Contributions

Conceptualization, R.Z.Y.; methodology, R.Z.Y.; software, R.Z.Y.; validation, R.Z.Y., D.D.W, and D.H.L.; formal analysis, R.Z.Y.; investigation, R.Z.Y.; resources, J.S.K., P.P.L. and S.A.L.; data curation, R.Z.Y.; writing—original draft preparation, R.Z.Y.; writing—review and editing, R.Z.Y.; visualization, R.Z.Y.; supervision, J.S.K.; project administration, J.S.K.; funding acquisition, R.Z.Y., J.S.K., P.P.L. and S.A.L.

## Animal Ethics

The study was conducted according to the guidelines of the Institutional Animal Care and Use Committee of the Zhengzhou University under project ID 2024-KY-0399-001, April 2024.

## Data Availability Statement

Data are available upon request from the authors.

## Reference

1. D. M. Cooper, N. Mons, J. W. Karpen, Adenylyl cyclases and the interaction between calcium and cAMP signalling. Nature. 374, 421–424 (1995).

2. C. Boularan, C. Gales, Cardiac cAMP: production, hydrolysis, modulation and detection. Front Pharmacol. 6, 203 (2015).

3. W. G. Robichaux, X. Cheng, Intracellular cAMP Sensor EPAC: Physiology, Pathophysiology, and Therapeutics Development. Physiological Reviews. 98, 919–1053 (2018).

4. E. E. Benarroch, HCN channels: function and clinical implications. Neurology. 80, 304–310 (2013).

5. S. Fenske, K. Hennis, R. D. Rötzer, V. F. Brox, E. Becirovic, A. Scharr, C. Gruner, T. Ziegler, V. Mehlfeld, J. Brennan, I. R. Efimov, A. G. Pauža, M. Moser, C. T. Wotjak, C. Kupatt, R. Gönner, R. Zhang, H. Zhang, X. Zong, M. Biel, C. Wahl-Schott, cAMP-dependent regulation of HCN4 controls the tonic entrainment process in sinoatrial node pacemaker cells. Nature Communications. 11, 5555 (2020).

6. S. Delhaye, B. Bardoni, Role of phosphodiesterases in the pathophysiology of neurodevelopmental disorders. Molecular Psychiatry. 26, 4570–4582 (2021).

7. R. X. Yamada, N. Matsuki, Y. Ikegaya, cAMP Differentially Regulates Axonal and Dendritic Development of Dentate Granule Cells*. Journal of Biological Chemistry. 280, 38020–38028 (2005).

8. D. Ohadi, D. L. Schmitt, B. Calabrese, S. Halpain, J. Zhang, P. Rangamani, Computational Modeling Reveals Frequency Modulation of Calcium-cAMP/PKA Pathway in Dendritic Spines. Biophys J. 117, 1963–1980 (2019).

9. T. F. Musial, E. Molina-Campos, L. A. Bean, N. Ybarra, R. Borenstein, M. L. Russo, E. W. Buss, D. Justus, K. M. Neuman, G. D. Ayala, S. A. Mullen, Y. Voskobiynyk, C. T. Tulisiak, J. A. Fels, N. J. Corbett, G. Carballo, C. D. Kennedy, J. Popovic, J. Ramos-Franco, M. Fill, M. R. Pergande, J. A. Borgia, G. T. Corbett, K. Pahan, Y. Han, D. M. Chetkovich, R. J. Vassar, R. W. Byrne, M. Matthew Oh, T. R. Stoub, S. Remy, J. F. Disterhoft, D. A. Nicholson, Store depletion-induced h-channel plasticity rescues a channelopathy linked to Alzheimer’s disease. Neurobiol Learn Mem. 154, 141–157 (2018).

10. X. Chang, J. Wang, H. Jiang, L. Shi, J. Xie, Hyperpolarization-Activated Cyclic Nucleotide-Gated Channels: An Emerging Role in Neurodegenerative Diseases. Frontiers in Molecular Neuroscience. 12 (2019), doi:10.3389/fnmol.2019.00141.

11. C. H. Good, A. F. Hoffman, B. J. Hoffer, V. I. Chefer, T. S. Shippenberg, C. M. Bäckman, N.-G. Larsson, L. Olson, S. Gellhaar, D. Galter, C. R. Lupica, Impaired nigrostriatal function precedes behavioral deficits in a genetic mitochondrial model of Parkinson’s disease. FASEB J. 25, 1333–1344 (2011).

12. L. E. Bleakley, R. J. Keenan, R. D. Graven, J. A. Metha, S. Ma, H. Daykin, L. Cornthwaite-Duncan, D. Hoyer, C. A. Reid, L. H. Jacobson, Altered EEG power spectrum, but not sleep-wake architecture, in HCN1 knockout mice. Behavioural Brain Research. 437, 114105 (2023).

13. C. Zhou, J. Liu, X.-D. Chen, General anesthesia mediated by effects on ion channels. World J Crit Care Med. 1, 80–93 (2012).

14. M. Shimizu, X. Mi, F. Toyoda, A. Kojima, W.-G. Ding, Y. Fukushima, M. Omatsu-Kanbe, H. Kitagawa, H. Matsuura, Propofol, an Anesthetic Agent, Inhibits HCN Channels through the Allosteric Modulation of the cAMP-Dependent Gating Mechanism. Biomolecules. 12 (2022), doi:10.3390/biom12040570.

15. X. Chang, J. Wang, H. Jiang, L. Shi, J. Xie, Hyperpolarization-Activated Cyclic Nucleotide-Gated Channels: An Emerging Role in Neurodegenerative Diseases. Front Mol Neurosci. 12, 141 (2019).

16. J. Hanoune, N. Defer, Regulation and role of adenylyl cyclase isoforms. Annu Rev Pharmacol Toxicol. 41, 145–174 (2001).

17. F. Zhang, L. Zhang, Y. Qi, H. Xu, Mitochondrial cAMP signaling. Cell Mol Life Sci. 73, 4577–4590 (2016).

18. R. D. Airan, K. R. Thompson, L. E. Fenno, H. Bernstein, K. Deisseroth, Temporally precise in vivo control of intracellular signalling. Nature. 458, 1025– 1029 (2009).

19. M. Iseki, S. Matsunaga, A. Murakami, K. Ohno, K. Shiga, K. Yoshida, M. Sugai, T. Takahashi, T. Hori, M. Watanabe, A blue-light-activated adenylyl cyclase mediates photoavoidance in Euglena gracilis. Nature. 415, 1047–1051 (2002).

20. V. Jansen, L. Alvarez, M. Balbach, T. Strünker, P. Hegemann, U. B. Kaupp, D. Wachten, Controlling fertilization and cAMP signaling in sperm by optogenetics. Elife. 4 (2015), doi:10.7554/elife.05161.

21. C. C. Doucette, D. C. Nguyen, D. Barteselli, S. Blanchard, M. Pelletier, D. Kesharwani, E. Jachimowicz, S. Su, M. Karolak, A. C. Brown, Optogenetic activation of UCP1-dependent thermogenesis in brown adipocytes. iScience. 26, 106560 (2023).

22. S. Yang, O. M. Constantin, D. Sachidanandan, H. Hofmann, T. C. Kunz, V. Kozjak-Pavlovic, T. G. Oertner, G. Nagel, R. J. Kittel, C. E. Gee, S. Gao, PACmn for improved optogenetic control of intracellular cAMP. BMC Biology. 19, 227 (2021).

23. H. Odaka, S. Arai, T. Inoue, T. Kitaguchi, Genetically-encoded yellow fluorescent cAMP indicator with an expanded dynamic range for dual-color imaging. PLoS One. 9, e100252 (2014).

24. R. Fischmeister, L. R. Castro, A. Abi-Gerges, F. Rochais, J. Jurevicius, J. Leroy, G. Vandecasteele, Compartmentation of cyclic nucleotide signaling in the heart: the role of cyclic nucleotide phosphodiesterases. Circulation research. 99, 816–828 (2006).

25. M. Xin, J. Feng, Y. Hao, J. You, X. Wang, X. Yin, P. Shang, D. Ma, Cyclic adenosine monophosphate in acute ischemic stroke: some to update, more to explore. Journal of the Neurological Sciences. 413 (2020), doi:10.1016/j.jns.2020.116775.

26. T. T. Luyben, J. Rai, H. Li, J. Georgiou, A. Avila, M. Zhen, G. L. Collingridge, T. Tominaga, K. Okamoto, Optogenetic Manipulation of Postsynaptic cAMP Using a Novel Transgenic Mouse Line Enables Synaptic Plasticity and Enhances Depolarization Following Tetanic Stimulation in the Hippocampal Dentate Gyrus. Front Neural Circuits. 14, 24 (2020).

27. A. Koschinski, M. Zaccolo, Activation of PKA in cell requires higher concentration of cAMP than in vitro: implications for compartmentalization of cAMP signalling. Sci Rep. 7, 14090 (2017).

28. G. Murer, K. Tseng, V. Sinay, R. Peñalva, I. Armando, J. H. Pazo, Turning behavior induced by injections of glutamate receptor antagonists into the substantia nigra of the rat. Synapse. 24, 147–155 (1996).

29. R. J. McPherson, J. F. Marshall, Substantia nigra glutamate antagonists produce contralateral turning and basal ganglia Fos expression: Interactions with D1 and D2 dopamine receptor agonists. Synapse. 36, 194–204 (2000).

30. G. Papadopoulos, J. P. Huston, Contralateral turning after unilateral electrolytic lesion of substantia nigra in thalamic rats. Neurosci Lett. 13, 63–67 (1979).

31. J. Scheel-Krüger, J. Arnt, G. Magelund, Behavioural stimulation induced by muscimol and other GABA agonists injected into the substantia nigra. Neuroscience Letters. 4, 351–356 (1977).

32. G. E. Martin, N. L. Papp, C. B. Bacino, Contralateral turning evoked by the intranigral microinjection of muscimol and other GABA agonists. Brain Research. 155, 297–312 (1978).

33. C. S. Chan, J. N. Guzman, E. Ilijic, J. N. Mercer, C. Rick, T. Tkatch, G. E. Meredith, D. J. Surmeier, “Rejuvenation” protects neurons in mouse models of Parkinson’s disease. Nature. 447, 1081–1086 (2007).

34. C. Carbone, A. Costa, G. Provensi, G. Mannaioni, A. Masi, The Hyperpolarization-Activated Current Determines Synaptic Excitability, Calcium Activity and Specific Viability of Substantia Nigra Dopaminergic Neurons. Front Cell Neurosci. 11, 187 (2017).

35. B. Hu, Q. Shi, Y. Guo, X. Diao, H. Guo, J. Zhang, L. Yu, H. Dai, L. Chen, The oscillatory boundary conditions of different frequency bands in Parkinson’s disease. J Theor Biol. 451, 67–79 (2018).

36. J. M. Shine, A. M. A. Handojoseno, T. N. Nguyen, Y. Tran, S. L. Naismith, H. Nguyen, S. J. G. Lewis, Abnormal patterns of theta frequency oscillations during the temporal evolution of freezing of gait in Parkinson’s disease. Clinical Neurophysiology. 125, 569–576 (2014).

37. W.-N. Xue, Y. Wang, S.-M. He, X.-L. Wang, J.-L. Zhu, G.-D. Gao, SK- and h-current contribute to the generation of theta-like resonance of rat substantia nigra pars compacta dopaminergic neurons at hyperpolarized membrane potentials. Brain Struct Funct. 217, 379–394 (2012).

38. A. G. Yee, S.-M. Lee, M. R. Hunter, M. Glass, P. S. Freestone, J. Lipski, Effects of the Parkinsonian toxin MPP+ on electrophysiological properties of nigral dopaminergic neurons. NeuroToxicology. 45, 1–11 (2014).

39. C. S. Chan, K. E. Glajch, T. S. Gertler, J. N. Guzman, J. N. Mercer, A. S. Lewis, A. B. Goldberg, T. Tkatch, R. Shigemoto, S. M. Fleming, D. M. Chetkovich, P. Osten, H. Kita, D. J. Surmeier, HCN channelopathy in external globus pallidus neurons in models of Parkinson’s disease. Nat Neurosci. 14, 85–92 (2011).

40. B. H. Meurers, G. Dziewczapolski, A. Bittner, T. Shi, F. Kamme, C. W. Shults, Dopamine depletion induced up-regulation of HCN3 enhances rebound excitability of basal ganglia output neurons. Neurobiol Dis. 34, 178–188 (2009).

41. A. Mathis, P. Mamidanna, K. M. Cury, T. Abe, V. N. Murthy, M. W. Mathis, M. Bethge, DeepLabCut: markerless pose estimation of user-defined body parts with deep learning. Nature Neuroscience. 21, 1281–1289 (2018).

